# The FANCD2-FANCI heterodimer coordinates chromatin openness and cell cycle progression throughout DNA double-strand break repair

**DOI:** 10.1101/2025.04.15.648866

**Authors:** Christine M. Joyce, Julien Bacal, Soham P. Chowdhury, Andrew N. Brown, Amy K. Wang, Carmen Cruz, Kameron Bains, Zachary N. Rodriguez, Nathan J. McCormick, Yaara Tzadikario, Katherine U. Tavasoli, Brooke M. Gardner, Chris D. Richardson

## Abstract

The FANCD2-FANCI heterodimer contributes to DNA repair at interstrand crosslinks and sites of replication stress. This complex has been physically and mechanistically linked to double-strand break (DSB) repair, but its role in that process remains undefined. Here we show that the FANCD2-FANCI heterodimer dynamically interacts with open chromatin regions, including transient, DSB-induced open chromatin, where it can be stabilized by co-activation by the DNA repair kinase ATM and the Fanconi anemia core ubiquitin ligase. The loaded FANCD2-FANCI heterodimer stabilizes open chromatin and promotes resection and loading of RPA through increased association of BRCA1 and BLM. Chromatin-loaded FANCD2-FANCI has a second distinct function promoting a G2 arrest that is dependent on the ATR-CHK1-WEE1 axis. Our results support a two-step genome surveillance model in which FANCD2-FANCI monitors open chromatin sites and is stably loaded to coordinate DNA repair activities in response to signaling from a DNA repair kinase.

## Introduction

Fanconi anemia (FA) is a genetic disorder characterized by developmental abnormalities, bone marrow failure in early childhood, and an increased incidence of cancer, primarily hematologic and solid tumors^1^. FA is caused by biallelic mutations in any of at least 22 genes that comprise the FA-BRCA pathway^2^. The defining characteristic of an FA gene is that FA cells undergo chromosomal breakage upon exposure to DNA crosslinking agents like mitomycin C.^1^ This test illustrates that the FA-BRCA pathway functions to repair DNA, but hints that the FA-BRCA pathway may unite multiple different repair activities to deal with complex lesions like interstrand crosslinks (ICLs). It remains unclear if the FA-BRCA pathway is activated by distinct DNA damage signals to perform lesion-specific repair or if there is a unifying model that describes the recruitment and activity of the pathway.

The key activation event in the FA-BRCA pathway is the clamping of the FANCD2-FANCI heterodimer onto DNA, facilitated by the monoubiquitination of FANCD2 and FANCI. Several FA genes (FANCA, B, C, E, F, G, L, Y along with FAAP20) encode proteins that form the FA core ubiquitin ligase complex, which monoubiquitinates FANCD2-FANCI^2^. Appearance of monoubiquitinated products on immunoblots or accumulation of ubiquitinated FANCD2-FANCI on chromatin is therefore used as a marker for underlying DNA damage^3^. Another FA factor, FANCM, likely recruits the core complex to sites of DNA damage^4,5^. The remaining genes in the FA-BRCA pathway include effector proteins that have well-defined roles in DNA repair, such as three RAD51 orthologs (FANCO/R/U), BRCA complex members (FANCD1/S/J/N), and nucleases (SLX4/XPF - FANCP/Q). The overall model that emerges from interactions within the FA-BRCA pathway indicates that DNA damage is sensed by the FA core complex, which clamps FANCD2-FANCI onto chromatin, wherein the heterodimer can facilitate downstream repair events. The function of individual FA genes reinforces the idea that it is a modular pathway that can be activated by multiple types of DNA damage and coordinate distinct repair activities.

The FA-BRCA pathway is best understood in its role in replication-dependent interstrand crosslink (ICL) repair. During this process, converging replication forks stall near the ICL, and FANCD2-FANCI associates with the lesion. The FA E3 ligase core complex monoubiquitinates the FANCD2-FANCI heterodimer at ICL-damaged chromatin. Subsequent recruitment of downstream factors excises crosslinks from DNA, and rescues severed replication forks^2^. This latter activity is similar to DSB repair. DSB repair proteins belong to one of two pathways: end-joining (EJ) proteins that reseal the lesion, or homologous recombination (HR) proteins that copy the sequence around the lesion from an intact template molecule^6^. HR requires the conversion of the DSB into a recombination filament, utilizing the nuclease activity of the MRE11-RAD50-NBS1 (MRN) complex. Next, activation of long-range 5’-3’ exonucleases BLM-DNA2 or EXO1 exposes single-stranded DNA, which RPA binds and protects from degradation by other nucleases. Finally, RAD51 replaces RPA, finalizing the formation of the recombination filament. BRCA1 and BRCA2 regulate events in this cascade, likely loading of RPA and RAD51, respectively. Outside of its canonical role in ICL repair, the FA-BRCA pathway has been shown to participate in fork protection and restart^7,8^, colocalize with R-loops^9^, maintain genome stability at chromosomal fragile sites^10^, participate in the repair of UV-induced lesions^11^, and localize to DSBs^12,13^. It has been challenging to build lesion-specific models for the activation and function of the FA-BRCA pathway due to the diverse range of DNA substrates.

*In vitro* characterization of the FA-BRCA pathway suggests a lack of lesion-specific recruitment and activation. DNA binding experiments revealed that FANCD2-FANCI binds to many naked DNA substrates *in vitro*, with the highest affinity for branched DNA substrates that resemble stalled replication forks^14^. Ubiquitination of FANCD2 by the FA core complex locks the closed FANCD2-FANCI heterodimer onto DNA substrates. High resolution cryo-EM structures show unmodified as well as monoubiquitinated forms of the FANCD2-FANCI heterodimer bound to double-stranded DNA^15–17^. These *in vitro* experiments describe a model in which the FANCD2-FANCI heterodimer binds to many DNA topologies and is then ubiquitinated by the FA core complex, undergoing a conformational change that increases binding affinity.

Our investigation into the recruitment and function of FANCD2-FANCI focused on its role at DSBs. Many FA-BRCA members, including FANCD2-FANCI, are genetically required for DSB repair, and depletion of these factors alters DSB repair outcomes^12^. FANCD2-FANCI is recruited to DSBs after early DSB repair complexes, including the EJ complex DNA-PK, but before factors like RAD51 that participate in HR^18^. These data support a model in which FA-BRCA members enable HR repair pathways. However, the precise role of the FANCD2-FANCI heterodimer at DSBs remains unclear. For instance, although FANCD2-FANCI associates with DSBs during repair, where we would expect to see FANCD2 bound to cut DNA, the majority of the DNA bound to FANCD2 does not end at the DSB, but in fact spans the DSB^19^. This observation is consistent with FANCD2-FANCI bound to DSBs that have already been repaired, and therefore appears incompatible with the timing of its arrival at chromatin. We therefore set out to understand the substrate that FANCD2-FANCI binds to and its role at DSBs.

We isolated the DNA bound by the FANCD2-FANCI heterodimer to determine its DNA substrate in the context of DSB repair. We show that the FANCD2-FANCI heterodimer interacts dynamically with stable open chromatin sites in human cells and with transient open chromatin sites created by DNA repair events at DSBs. We use chromosome location- and cell cycle-specific assays to study FANCD2-FANCI at DSBs separate from their other replication or genome-maintenance functions. We find that FANCD2-FANCI is loaded onto chromatin in response to signaling at DSBs. Additionally, we find that FANCD2-FANCI is an integral part of the DNA resection machinery that assembles at DSBs, promoting resection and RPA loading. Finally, we show that FANCD2 alters cell cycle progression after DSB induction by promoting a G2 arrest via the ATR-CHK1-WEE1 pathway. Overall, our results support a model in which FANCD2-FANCI monitors open chromatin sites and regulates the accessibility of these sites to DNA repair proteins.

## Results

### FANCD2-FANCI binds both single- and double-stranded DNA at DSBs, remaining bound throughout DNA resection

Past studies have shown that FANCD2 is robustly recruited to DSBs, but the DNA bound to FANCD2 in these experiments was primarily uncut^19^. This observation is difficult to reconcile with other results showing that disruption of FANCD2 alters DSB repair outcomes^20^. To refine our understanding of FANCD2-FANCI interaction with damaged DNA, we performed inducible DSB assays at single or multiple loci. We monitored the induction of DSBs at 122 sites using the AsiSI endonuclease (5’-GCGATCGC-3’), 50 sites using the AluGG guide complexed with Cas9, or single sites using HBB, LMNB1, RAB11A, or HIST1H2BJ guides complexed with Cas9 (**Figure 1A, Table S1**)^21–23^. After DSB induction, we performed ChIP-seq experiments that preserved DNA strand information, and thus measured resected DNA (**Figure 1B**).

**Figure 1:**
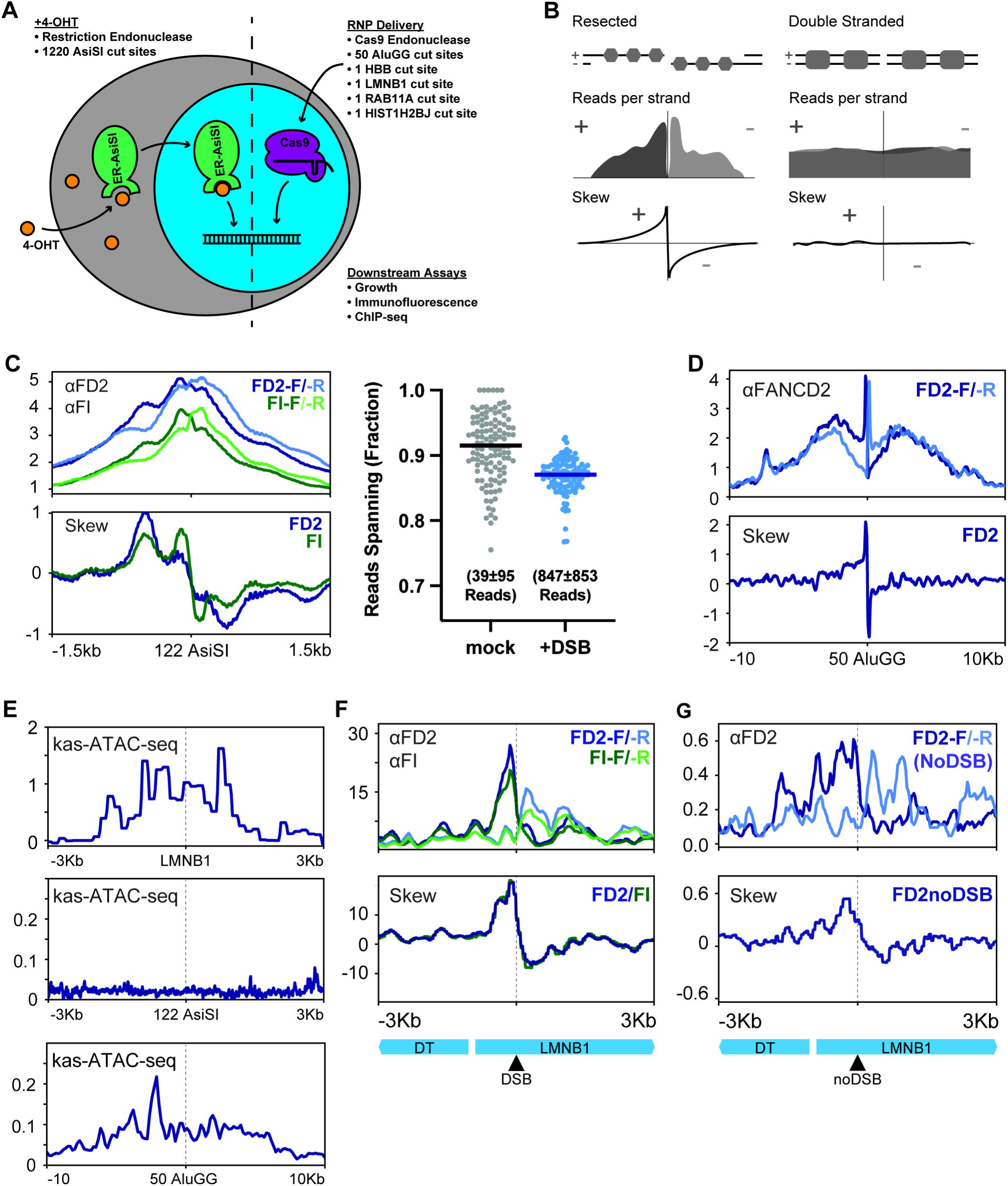
The FANCD2-FANCI heterodimer binds double- and single-stranded DNA at DSB sites. A. DSB induction schematic showing: 1) Addition of 4-OHT driving nuclear localization of the ER-AsiSI fusion protein and digestion of target sites; or 2) electroporation of Cas9 and guide RNA targeting single or multiple genomic loci. B. Stranded ChIP-seq schematic showing resected or double-stranded DNA, read profiles consistent with resected or double-stranded DNA, and skew profiles generated by subtracting reads mapping to the – strand from reads mapping to the + strand. C. Stranded ChIP-seq data presented as reads mapping to distinct strands (top) or skew (bottom) at 122 AsiSI sites 4 hours after DSB induction. Individual reads in FANCD2 ChIP datasets spanning the DSB site were plotted as a frequency (right). Immunoprecipitations were performed using FANCD2 or FANCI antibodies as indicated. Line plots are representative of n=2 biological replicates and dot plots present all data from n=2 replicates in each condition. D. Stranded ChIP-seq data presented as reads mapping to distinct strands (top) or skew (bottom) at 50 AluGG sites 16 hours after DSB induction. Immunoprecipitations were performed using a FANCD2 antibody. All plots are representative of n=2 biological replicates. E. Kas-ATAC-seq in GM12878 cells reproduced from PRJNA1103095 for the LMNB1 site (top), 122 AsiSI sites (middle), or 50 AluGG sites (bottom). All plots are the average of n=2 biological replicates. F. Stranded ChIP-seq data presented as reads mapping to distinct strands (top) or skew (bottom) at the LMNB1 cut site 16 hours after DSB induction. Immunoprecipitations were performed using FANCD2 or FANCI antibodies as indicated. Line plots are representative of n=2 biological replicates. G. Stranded ChIP-seq data presented as reads mapping to distinct strands (top) or skew (bottom) at the LMNB1 cut site in its natural (no DSB) context. Line plots are representative of n=2 biological replicates.

We observed that FANCD2-FANCI binds to DSBs 4 hours after AsiSI induction (**Figure 1C** – top left plot). Consistent with previous results at single DSBs^19^, most of the reads spanned the cut site, indicating that FANCD2-FANCI is mostly bound to intact DNA (**Figure 1C** – right plot). FANCD2-FANCI binding at AsiSI DSBs on average showed a preference for the 3’ strand on either side of the DSB, indicating that FANCD2-FANCI can bind single-stranded DNA. This trend can be seen by plotting read counts mapping to the + or – DNA strands, or alternatively by plotting read skews that report the excess of reads mapping to the + or – DNA strands on either side of the DSB (**Figure 1B, 1C** – bottom left plot). FANCD2 binds to AluGG DSB sites 16 hours after DSB induction with a wider distribution and greater preference for the 3’ strand on either side of the DSB (**Figure 1D**). The width and strand preference of FANCD2-FANCI distributions at AsiSI and AluGG sites are consistent with the timing of DSB maturation events. For example, extensive 5’ to 3’ resection, which leaves single-stranded 3’ ends, would occur from 4 to 16 hours after DSB formation. Overall, these results demonstrate that the FANCD2-FANCI heterodimer retains its binding during transitions at the DSB, binding both double-stranded DNA prior to resection activity and single-stranded DNA after resection activity.

Altogether, these data suggest that FANCD2 can remain bound throughout the repair process – even beyond resealing of the DSB. For example, DSB induction using a specific guide RNA that targets the N-terminus of the HBB gene causes a 9bp deletion of sequence adjacent to the DSB site. Immunoprecipitation of FANCD2 from cells with DSBs induced at HBB purifies DNA containing this 9bp deletion (**Figure S1A**). This finding indicates that the FANCD2-FANCI heterodimer binds prior to DSB repair and remains bound throughout the repair process, including post-repair of a DSB.

To better understand FANCD2-FANCI association at an individual DSB, we focused on a gRNA targeting the N-terminus of the LMNB1 gene. The resulting DSB at LMNB1 occurs in a region where LMNB1 and LMNB1-DT are divergently transcribed, so we sought to determine whether the FANCD2-FANCI distribution at this cut site reflected an underlying chromatin state. To investigate the chromatin state at this locus, we turned to existing Kethoxal-Assisted Single-stranded DNA Assay for Transposase-Accessible Chromatin with Sequencing (kas-ATAC-seq) datasets^24^. We noted that the DNA in the LMNB1 region is single-stranded and accessible prior to any DSB induction (**Figure 1E** – top plot). AsiSI and AluGG sites did not show the same magnitude of kas-ATAC-seq signal, indicating that uncut regions containing the target sequence prior to the induction of DSBs are on average double-stranded (**Figure 1E** – middle and bottom plots, respectively). Strand-specific sequencing of DNA bound to FANCD2 or FANCI at the LMNB1 DSB revealed that the FANCD2-FANCI heterodimer is bound primarily to the 3’ terminated strand on either side of the break (**Figure 1F**). In comparison to the AsiSI and AluGG break sites, FANCD2-FANCI is asymmetrically bound at the LMNB1 break site, with more association towards the DT gene. Upon further investigation of this asymmetry, we observed by ChIP-seq a stranded and asymmetric FANCD2 binding pattern even in mock treated cells (no LMNB1 DSBs), although the magnitude of signal was 30-fold less without DSB induction (**Figure 1G**). This finding suggests that the FANCD2-FANCI heterodimer not only binds throughout the transitions of DSB repair but also binds prior to DNA damage.

Overall, these results demonstrate that FANCD2-FANCI binds to single-stranded DNA present before the DSB is introduced, as is the case with LMNB1, or single-stranded DNA created by DSB resection, as is the case for AsiSI and AluGG DSBs. More broadly, this finding indicates that the FANCD2-FANCI heterodimer retains its binding distribution during transitions, either from double-stranded DNA to single-stranded DNA during resection, or from a pre-DSB to post-DSB chromatin state.

### FANCD2-FANCI is preloaded at open chromatin sites and further accumulates at DSB-induced open chromatin

The existence of a pre-DSB distribution of FANCD2 at LMNB1 led us to investigate if chromatin-bound FANCD2-FANCI in uncut ChIP-seq datasets reflects a broad affinity for open chromatin.

To assess whether FANCD2-FANCI binds at sites of open chromatin, we compared FANCD2-FANCI binding before and after DSB induction at many sites in the genome using AsiSI endonuclease. Induction of DSBs by addition of 4-OHT to cultures for 4 hours resulted in accumulation of FANCD2 at DSB sites (**Figure 2A** – third plot). We verified that these sites were cut by also monitoring recruitment of the DSB-repair factor, MRE11 (**Figure 2A** – fourth plot). However, many of the AsiSI sites that accumulate MRE11 and FANCD2 are also bound by FANCD2 prior to DSB induction (**Figure 2A** – second plot). To determine whether these 122 AsiSI sites overlap with open chromatin, we performed ATAC-seq to identify regions of open chromatin (**Figure 2A** – first plot). Spearman correlations between these datasets indicate that chromatin openness correlates with loading of FANCD2 in the absence of DSBs (**Figure 2B**). Furthermore, these correlations strongly predict the magnitude of FANCD2 association when DSBs are induced. We thus conclude that AsiSI sites in our dataset overlap with open chromatin and have underlying FANCD2-FANCI distributions, which are reinforced upon DSB induction^25^.

**Figure 2:**
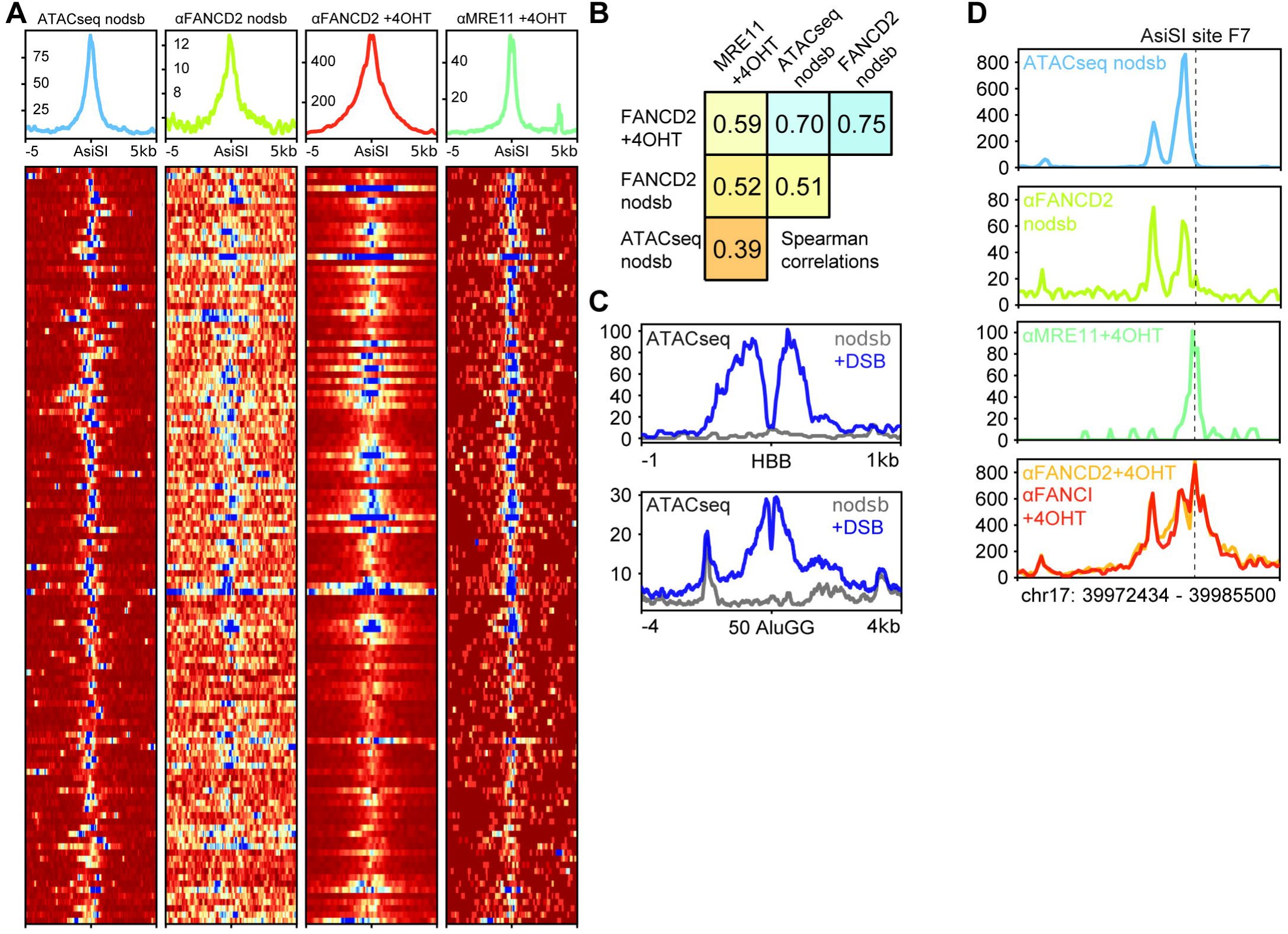
The FANCD2-FANCI heterodimer interacts dynamically with induced and stable open chromatin sites in human cells. A. ATAC-seq and ChIP-seq data presented as average and individual signal at 122 AsiSI sites either uncut (nodsb) or 4 hours after DSB induction (+4OHT) and showing chromatin openness or repair factor recruitment. Immunoprecipitations were performed using MRE11 or FANCD2 antibodies as indicated. All plots are representative of n=2 biological replicates. B. Strong (≥0.7), moderate (≥0.5), and weak (<0.5) Spearman correlations between signal at all (1228) AsiSI consensus sites generated by ATAC-seq without DSBs, FANCD2 without DSBs, FANCD2 with DSBs, and MRE11 with DSBs. Data was generated from n=2 biological replicates for each condition. C. ATAC-seq signal at single (HBB) or multiple (50 AluGG) DSB sites without (nodsb) or 4 hours after DSB induction (+DSB). Each panel shown is n=1 biological replicate. D. ATAC-seq and ChIP-seq data presented as signal at the F7 AsiSI site showing open chromatin (top), recruitment of FANCD2 without DSBs (second from top), recruitment of MRE11 with DSBs (second from bottom), and recruitment of FANCD2/FANCI to DSBs (bottom). Data shown are representative of n=2 biological replicates.

In contrast, Alu elements tend to occur in chromatin that is inaccessible to ATAC-seq. In the absence of DSBs, we see low levels of FANCD2 binding to 50 AluGG sites (**Figure S1B**). Upon DSB induction, FANCD2 robustly accumulates at these sites, binding both forward and reverse strands, as well as DNA that spans the cut site (**Figure 1D, S1C**). The FANCD2 distribution at AluGG DSB sites is biased towards binding to the 3’-terminated strand, consistent with 5’ resection at the DSB (**Figure 1D**). Additionally, we find that FANCD2-FANCI is bound to a lower fraction of DNA spanning AluGG DSB sites, consistent with non-DSB FANCD2 binding contributing to the population of ChIP reads spanning the DSB site (**Figure S1C**).

Overall, these results support a model in which FANCD2-FANCI binds DNA with a background, constitutive profile that follows the underlying chromatin accessibility. When FANCD2-FANCI is preloaded near a region to be targeted for a DSB, the background binding pattern is preserved and amplified by the formation of the adjacent DSB. DSBs themselves cause transient open chromatin (**Figure 2C**)^22^, which indicates that recruitment of FANCD2-FANCI to DSBs is chromatin-directed. An example of this activity can be seen at AsiSI site F7, which has three open chromatin regions that associate with FANCD2 in the absence of DSBs, but do not overlap the consensus cut site (**Figure 2D**). In the presence of DSBs, FANCD2 associates with both the DSB and the adjacent open chromatin sites; the overall amount of FANCD2 ChIP-seq signal increases ∼10-fold after DSB induction. We therefore hypothesize that open chromatin – not DNA lesions – is the main substrate for FANCD2-FANCI.

### Binding of FANCD2-FANCI to open chromatin substrates requires the FA core complex

To further test if FANCD2-FANCI binds open chromatin substrates, we compared both FANCD2 and FANCI binding to non-TSS DNase hypersensitive sites defined by the ENCODE consortium (**Table S1**) in osteosarcoma MG63 cells (**Figure 3A**)^26^. We found that FANCD2-FANCI is enriched at non-TSS DNase hypersensitive sites in the absence of DSBs. In addition to genomic open chromatin regions, our ChIP-seq datasets also revealed that FANCD2-FANCI bound to non-genomic open chromatin regions, including mitochondrial DNA^27^ and extrachromosomal DNA (**Figure S2A**)^28^.

**Figure 3:**
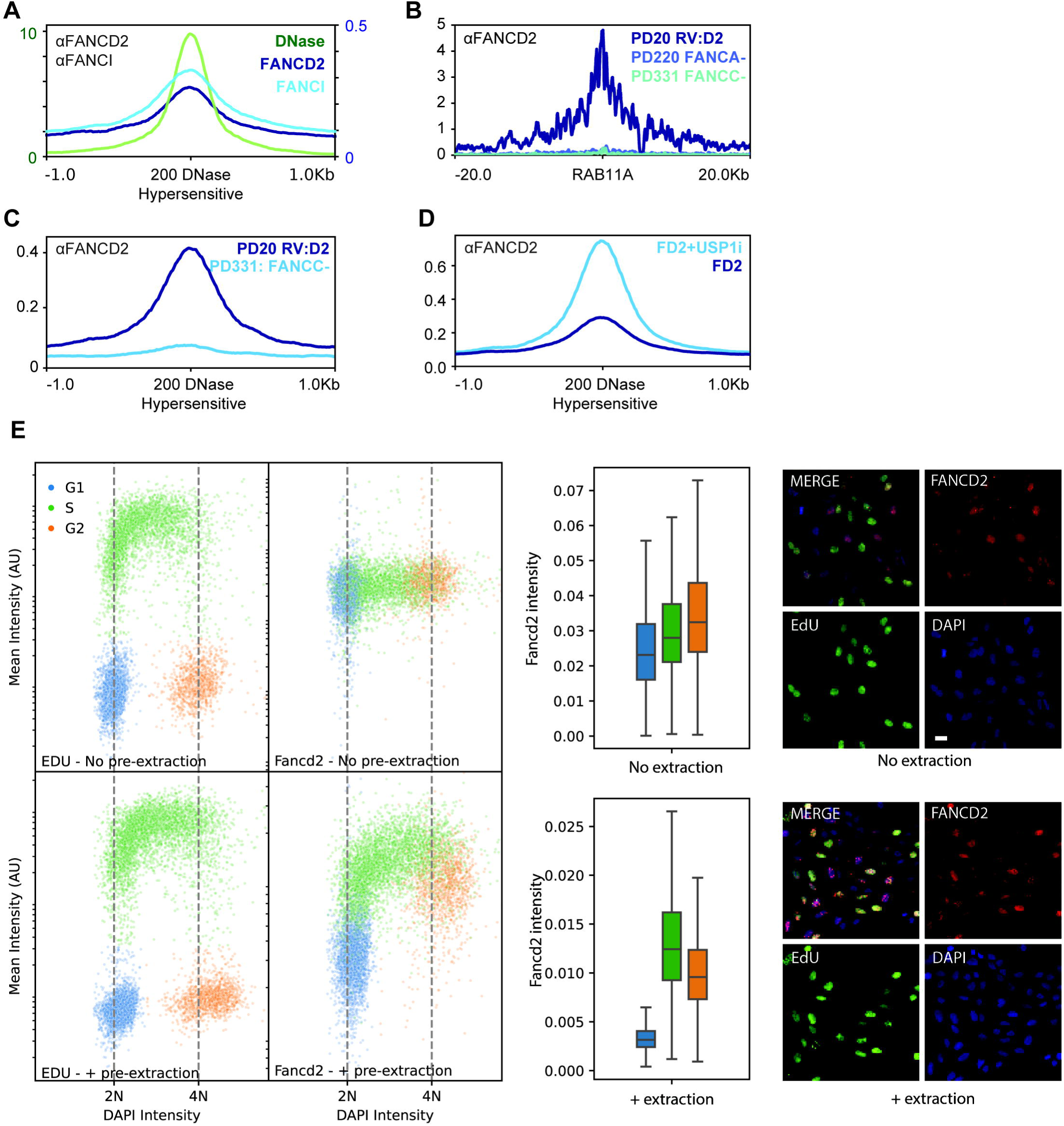
FANCD2-FANCI loading at open chromatin substrates requires the FA core complex and is cell cycle dependent. A. ChIP-seq and DNase hypersensitivity profiles showing FANCD2 binding, FANCI binding, and DNase hypersensitivity at 200 non-TSS DNase hypersensitive sites. Data shown are representative of n=2 biological replicates. B. ChIP-seq data showing FANCD2 binding to a DSB in the RAB11A gene in cell lines expressing FANCD2 (PD20 RV:D2), or lacking the FA core components FANCA (PD220) or FANCC (PD331). Data shown are representative of n=2 biological replicates. C. ChIP-seq profiles showing FANCD2 binding to DNase hypersensitive sites in asynchronous cells with and without the core complex component FANCC. Data shown are representative of n=2 biological replicates. D. ChIP-seq profiles showing FANCD2 binding to DNase hypersensitive sites in asynchronous cells with and without inhibition of the deubiquitinase USP1. Data shown are representative of n=2 biological replicates. E. QIBC immunofluorescence tracking EdU incorporation, DAPI incorporation, and FANCD2 chromatin binding in unextracted (top row) or pre-extracted cells (bottom row). Data are presented as pseudo-cell cycle plots (left), FANCD2 mean intensity as a function of cell cycle (middle), and example IF images (right). Data shown are generated from at least m=12 images and n=2000 cells. Scale bar = 20μm.

Consistent with past studies, we observed that FANCD2 is robustly recruited to DSBs in an FA-core dependent manner (**Figure 3B**). Therefore, we sought to determine whether the break independent binding of FANCD2-FANCI to open chromatin substrates also requires the FA core complex. In FANCC null cell lines, FANCD2 abundance decreased at DNase hypersensitive sites, indicating that the binding of FANCD2-FANCI to these sites is stabilized by the FA core complex (**Figure 3C**). Additionally, we found that binding to plasmid-derived extrachromosomal DNA depends on the FA core (**Figure S2B**). Conversely, we tested the effect of inhibition of the FANCD2-FANCI deubiquitinase, USP1. Treatment of cells with a USP1 inhibitor^29^, which prevents de-ubiquitination of FANCD2-FANCI, increased binding of FANCD2 at DNase hypersensitive sites (**Figure 3D**). USP1 inhibition also reduced recombination between a genomic DSB and genomic template provided by the DR-GFP system (**Figure S2C**)^30^. This observation is consistent with overloading of FANCD2-FANCI at non-DSB sites, thereby depleting the cellular pool of FANCD2-FANCI that is available for DNA repair. Together, these results support a model in which core-dependent FANCD2-FANCI loading occurs at open chromatin substrates.

### Binding of FANCD2-FANCI is cell cycle dependent

To understand if FANCD2-FANCI binding to open chromatin changes throughout the cell cycle, we performed cell cycle resolved immunofluorescence experiments using Quantitative Image Based Cytometry (QIBC)^31^. We discovered that FANCD2-FANCI is bound to chromatin throughout the S and G2 phases, unloaded from chromatin in mitosis before G1 entry, and excluded from chromatin in G1 (**Figure 3E** – plus extraction). These changes are driven by chromatin loading and not protein abundance, as FANCD2 expression does not vary dramatically throughout the cell cycle (**Figure 3E** – no extraction). To understand how cells establish FANCD2-FANCI binding during S phase, we separated regions of the genome into early and late replicating clusters based on REPLIseq datasets^32^, and we examined the binding of FANCD2 to DSB sites or open chromatin sites in these regions. Replication status (early/late) did not influence FANCD2 localization to AsiSI DSBs (**Figure S2D** – top plots). However, there was an ∼2-fold increase in FANCD2 signal at ATAC-seq sites located in early replicating regions relative to late replicating regions of the genome (**Figure S2D** – bottom plots). We conclude that replication does not play a role in recruiting FANCD2 to DSBs. However, replication may play a minor role in recruiting FANCD2 to open chromatin sites. Overall, we favor a model in which FANCD2-FANCI samples open chromatin in the S and G2 phases of the cell cycle, and therefore is not directly targeted to its substrates by replication forks. This model is further supported by the binding of FANCD2 to plasmid-derived extrachromosomal DNA, which does not replicate in human cells yet loads FANCD2 with high abundance (**Figure S2A**).

### ATM activation stabilizes FANCD2-FANCI at open chromatin sites during resection in the vicinity of DSBs

To establish the timing of recruitment after DSB induction, we monitored the association of FANCD2 and the DSB-binding protein MRE11 with Cas9-induced DSBs in the RAB11A promoter, a region of open chromatin. We found that MRE11 association peaks approximately 8 hours after DSB induction (**Figure 4A** – left plot). FANCD2 slowly accumulated after DSB induction and continued to accumulate out to 24 hours. The majority of the DNA bound to MRE11 ended at the cut site, indicating MRE11 is bound to unrepaired DNA (**Figure 4A** – right plot). FANCD2 on the other hand is primarily bound to DNA that spans the cut site, consistent with a fraction of FANCD2 preloaded at open chromatin at the RAB11A locus. To determine if this order is coincidental or an indication of direct mechanistic linkage, we inhibited MRE11 using the exonuclease domain inhibitor Mirin or the endonuclease domain inhibitor PFM01^33^. Inhibition of either nuclease domain of MRE11 prevented FANCD2 binding, with endonuclease inhibition causing the greatest reduction in FANCD2 binding at AsiSI-induced DSBs (**Figure 4B** – top plot). Neither inhibitor affected FANCD2 binding to non-DSB open chromatin sites (**Figure 4B** – bottom plot). We conclude that FANCD2 accumulation occurs primarily after activation of the nuclease domains of MRE11, which contribute to short-range resection activity. These results are consistent with FANCD2 binding to single-stranded DNA within open chromatin after DNA resection is initiated at DSBs.

**Figure 4:**
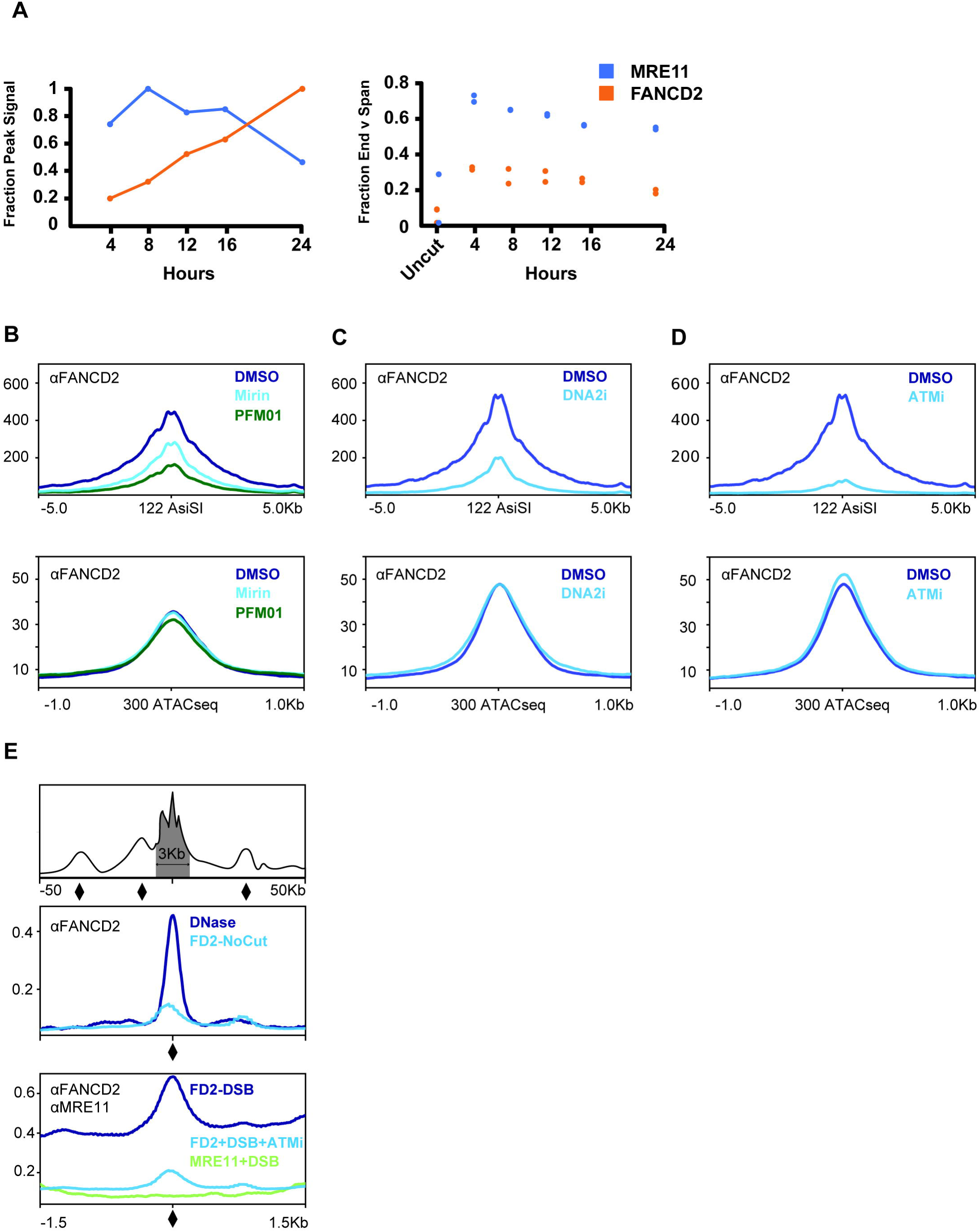
ATM activation stabilizes FANCD2-FANCI at DSBs and DSB-adjacent DNase hypersensitive sites. A. MRE11 and FANCD2 binding as a function of time after DSB induction at RAB11A (left panel) and the fraction of reads ending at the DSB sites in each condition (right panel). Data shown are generated from n=2 biological replicates. B. ChIP-seq summary data showing FANCD2 binding to DSBs (top) or open chromatin sites (bottom) in untreated, mirin-treated, or PFM01-treated conditions. Data shown are representative of n=2 biological replicates. C. ChIP-seq summary data showing FANCD2 binding to DSBs (top) or open chromatin sites (bottom) in untreated, or NSC-105808-treated conditions. Data shown are representative of n=2 biological replicates. D. ChIP-seq summary data showing FANCD2 binding to DSBs (top) or open chromatin sites (bottom) in untreated, or KU-55933-treated conditions. Data shown are representative of n=2 biological replicates. E. ChIP-seq and DNase-hypersensitive data showing FANCD2 or MRE11 association at DSB-adjacent DNase-hypersensitive sites. These sites arise more than 1.5kb but less than 50kb upstream and downstream of 122 AsiSI cut sites (top panel) and average signal at these sites is presented without (middle panel) or with (bottom panel) DSB induction. Data shown are representative of n=2 biological replicates.

MRE11 binding is upstream of additional events in DSB repair, most notably the activity of long-range exonucleases, such as DNA2, and the activation of the DNA repair signaling kinase, ATM. To understand how each of these events contribute to FANCD2 association with DSBs, we inhibited both long-range exonuclease activity and ATM activation. Inhibition of the long-range exonuclease DNA2 with NSC-105808 reduced FANCD2 association at DSBs, but not at non-DSB open chromatin sites (**Figure 4C**). NSC-105808 has previously been shown to reduce the accumulation of phosphorylated RPA (a readout for DNA resection) upon treatment of cells with the topoisomerase inhibitor camptothecin^34^. We therefore interpret the reduction in FANCD2 loading at DSBs in the presence of PFM01, Mirin, and NSC-105808 as an overall reduction in the footprint of open chromatin surrounding the DSB in the absence of resection activity. The lack of change in binding at non-DSB open chromatin sites with these small molecule inhibitors further indicates that resection activity at DSBs is critical to altering the chromatin state that leads to the accumulation of FANCD2 at DSBs.

DSBs activate the DNA repair signaling kinase ATM, which is recruited to DSBs by MRE11. Inhibition of ATM with KU-55933^35,36^ reduced FANCD2 association at DSBs, but not at non-DSB open chromatin sites (**Figure 4D**). Inhibition of ATM also reduced FANCD2 association at DSB-adjacent open chromatin sites (**Figure 4E**). To investigate this, we compiled a list of DSB-adjacent open chromatin sites ranging from 1.5kb to 50kb upstream or downstream from DSBs (**Figure 4E** – indicated by the diamonds on the representative first plot). These sites are DNase hypersensitive and load FANCD2 in the absence of DSBs (**Figure 4E** – second plot). When DSBs are induced, these sites accumulate additional FANCD2, but not MRE11, indicating that they are DSB-adjacent open chromatin sites, but not themselves DSBs (**Figure 4E** – third plot). Treatment with ATM inhibitor prevents additional FANCD2 accumulation at these sites (**Figure 4E** – compare third plot with second plot). We thus conclude that ATM inhibition reduces FANCD2 association with both DSBs and local, but not global, sites of open chromatin, as we observed no change in FANCD2 association at global non-DSB open chromatin sites (**Figure 4D** – bottom plot). We speculate that ATM inhibition, which prevents the phosphorylation activity of ATM kinase, subsequently prevents ubiquitination of FANCD2-FANCI, which would clamp the heterodimer at DSBs. Consistent with this hypothesis, DSB-adjacent DNase hypersensitive sites, located 50kb upstream or downstream of the DSB, lie within the footprint of γ-H2AX surrounding AsiSI cut sites^37^, which is a known ATM phosphorylation substrate (**Figure S3A**).

ATR, another DNA repair signaling kinase, promotes association of FANCD2-FANCI with stalled replication forks^38^. However, ATR inhibition does not alter FANCD2 recruitment to DSBs 4 hours after DSB induction in asynchronously dividing cells (**Figure S3B**). A longer 24-hour incubation with ATR inhibitor dramatically reduces FANCD2 binding at DSBs (**Figure S3C** – compare right 24H plot with left 4H plot). As ATR depletion causes a G1 arrest (and FANCD2 does not bind chromatin in G1), any depletion of FANCD2 at this timepoint is likely secondary to cell cycle effects.

These results support an overall model for FANCD2-FANCI as a DNA surveillance complex. FANCD2-FANCI samples open chromatin structures that exist naturally in cells or are formed as the product of chromatin remodeling events, for instance that arise during DSB repair. Activity of a kinase, ATM in the context of DSBs, primes the FANCD2-FANCI heterodimer for ubiquitination and clamping onto chromatin. This model broadly suggests that multiple kinases can regulate the association of FANCD2-FANCI at open chromatin in a lesion-specific manner.

### FANCD2-FANCI loading changes the recruitment of DNA repair proteins and chromatin accessibility in the vicinity of DSBs

To understand how FANCD2 binding alters DSB repair, we first monitored chromatin accessibility in FANCD2-depleted cells after induction of a DSB with either HBB or AluGG targeted Cas9. Depletion of FANCD2 decreased ATAC-seq signal at the HBB locus and at the 50 AluGG loci (**Figure 5A**). This result indicates that one role of FANCD2 is promoting chromatin openness at lesions. Since FANCD2 binds open chromatin, we surmise that the stabilization of open chromatin could be due directly to FANCD2-FANCI binding, or due to its recruitment of other DNA repair factors.To test if FANCD2-FANCI depletion alters DSB maturation, we assessed the effect of depleting FANCD2 on events at AsiSI DSBs. We first utilized a previously established quantitative PCR assay to measure the amount of single-stranded DNA^39^. Depletion of FANCD2 using siRNA reduces the amount, but not the extent, of DSB resection at AsiSI sites (**Figure 5B**). We see a similar reduction in the binding of RPA, a single-stranded binding protein used as a proxy for DNA resection, in CRISPRi cells depleted of FANCD2 using ChIP-seq (**Figure 5C** – fourth plot). Depletion of FANCD2 also reduces the abundance of BRCA1 and BLM helicase at DSBs (**Figure 5C** – second and third plots, respectively). Separate investigation of RPA, BLM, and BRCA1 loading at an individual AsiSI DSB after siRNA depletion of FANCD2 using ChIP-qPCR confirms that these factors are indeed reduced (**Figure S4A**). We note that FANCD2-FANCI, BRCA1, BLM, and RPA share the same distribution at AsiSI DSBs, which suggests a model in which these factors work together during DSB repair (**Figure 5C** – compare the width of the distributions across all four plots).

**Figure 5:**
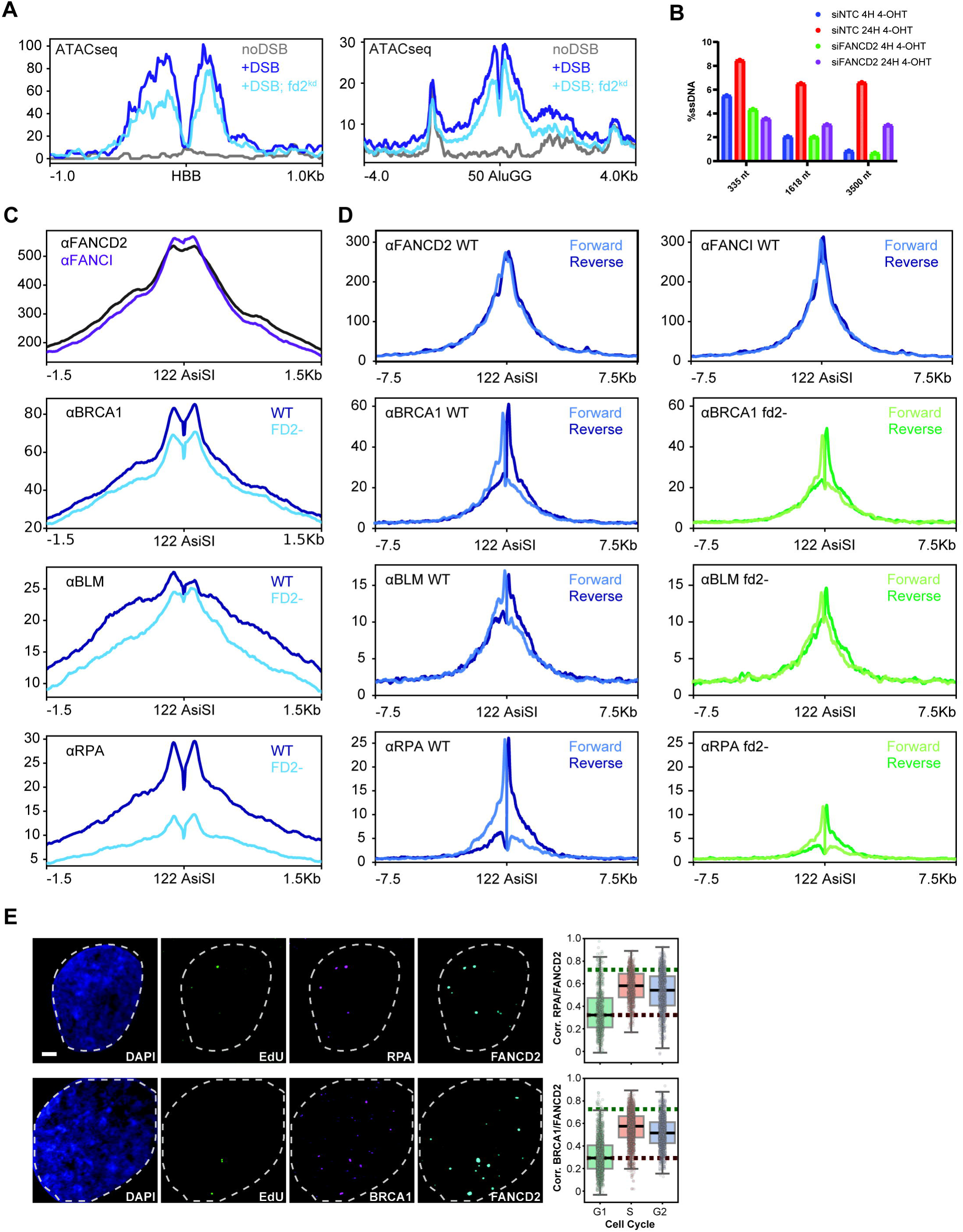
FANCD2-FANCI loading changes the recruitment of DNA repair proteins and chromatin factors in the vicinity of DSBs. A. Summary ATAC-seq data showing chromatin accessibility at HBB (left) or the average of 50 AluGG cut sites (right) in FANCD2 knockdown or non-targeting control cells without DSBs or 4 hours after DSB induction. Each panel shown is n=1 biological replicate. B. qPCR-based DSB resection assay showing ssDNA abundance at three different distances from the DSB in mock treated (siNTC) and FD2-depleted cells (siFANCD2) 4 and 24 hours after DSB induction. Data were generated from n=3 technical replicates. C. ChIP-seq data showing FANCD2 and FANCI, BRCA1, BLM, and RPA signal at AsiSI DSBs in CRISPRi non-targeting control (WT) and FANCD2 knockdown (fd2-) cells 4 hours after DSB induction. Data shown are representative of n=2 biological replicates. D. Stranded ChIP-seq data showing FANCD2 and FANCI, BRCA1, BLM, and RPA signal at AsiSI DSBs in CRISPRi non-targeting control (WT) and FANCD2 knockdown (fd2-) cells 4 hours after DSB induction. Data shown are representative of n=2 biological replicates. E. QIBC data showing colocalization of DAPI, EdU, and RPA/BRCA1 with FANCD2. Example images are presented at left and correlation is plotted at right. Data shown are generated from at least m=12 images and n=2000 cells. Scale bar = 2μm.

We then investigated the strandedness of the DNA bound to each of these repair factors in unperturbed and FANCD2-depleted cells. The FANCD2-FANCI heterodimer binds with a slight preference for the 3’ terminated strand on either side of the DSB (**Figure 5D** – top plots). BRCA1 and BLM are preferentially bound to the 3’ terminated strand (**Figure 5D** – middle plots). RPA, a known single-stranded binding complex, is bound primarily to the 3’ terminated strand (**Figure 5D** – bottom plots). Depletion of FANCD2 using CRISPRi alters the abundance of these factors, but not the strandedness of the underlying DNA bound by the factors (**Figure 5D**). This result is consistent with our observation that depletion of FANCD2 reduced the amount, but not the extent, of DSB resection (**Figure 5B**). Altogether, our results support a model in which FANCD2 acts during the end-resection phase of DSB maturation.

To assess if repair proteins are bound to the same DNA lesion as FANCD2 simultaneously, we used QIBC to assess colocalization in G2. We chose G2 because it is a point in the cell cycle when FANCD2 can be loaded onto chromatin (**Figure 3E**), but recruitment of DNA repair proteins to sites of replication-stress is minimal. Using this approach, we found that FANCD2 and RPA, as well as FANCD2 and BRCA1, colocalize at DSBs in G2 (**Figure 5E** – representative immunofluorescence images). We quantified aggregate overlap by correlating the signal in multiple channels (see **Materials and Methods**) and comparing that signal to conditions where no correlation can occur (FANCD2 does not load onto chromatin in G1) or where correlation always occurs (e.g. phosphorylated ATM and γ-H2AX – **Figure S4H**). Pairwise correlations between RPA and FANCD2 indicate that RPA and FANCD2 strongly colocalize in G2 (**Figure 5E** – top plot). Similarly, pairwise correlations between BRCA1 and FANCD2 indicate that BRCA1 and FANCD2 also strongly colocalize in G2 (**Figure 5E** – bottom plot).

The largest effects of FANCD2 depletion that we observed were on DNA resection and RPA abundance – and not the recruitment of BRCA1 or BLM. This supports a model in which the FANCD2-FANCI heterodimer acts alongside BRCA1 and BLM to facilitate DNA resection and load RPA onto resected DNA. This partial reduction in RPA loading at DSBs likely explains the decrease in recombination seen in FANCD2-deficient cell lines (**Figure S4B**). Consistent with FANCD2 working with BRCA1 to promote RPA loading, co-depletion of FANCD2 and BRCA1 reduces recombination frequencies in the context of genomic DSB and template provided by the DR-GFP system (**Figure S4B**).

We observed no decrease in MRE11 abundance at DSBs in FANCD2-depleted cells, consistent with its arrival at lesions before FANCD2 (**Figure S4C** – top plot). Strikingly, we also observed no decrease in BRCA2 abundance at DSBs in FANCD2-depleted cells (**Figure S4C** – bottom plot). Moreover, both MRE11 and BRCA2 share the same narrow distribution at AsiSI DSBs (**Figure S4C** – compare top plot with bottom plot). Further investigation of the strandedness of the DNA bound to each these repair factors in unperturbed and FANCD2-depleted cells revealed that both MRE11 and BRCA2 bind with a slight preference for the 3’ terminated strand on either side of the DSB (**Figure S4D**). Altogether, since FANCD2 depletion does not decrease BRCA2 loading at DSBs, and since BRCA2 has a narrow distribution, similar to MRE11, we hypothesize that BRCA2 may act relatively early in DSB repair, likely prior to resection activity.

FANCD2 depletion causes a modest reduction of RAD51 loading at DSBs (**Figure S4E**). After DNA resection, RPA is replaced by RAD51, in a process facilitated by BRCA2^6^. Further investigation of the strandedness of the DNA bound to RAD51 in unperturbed and FANCD2-depleted cells revealed that RAD51, like RPA, is bound primarily to the 3’ terminated strand (**Figure S4F**). We hypothesize that this result is likely the average of two opposing activities: decreased DNA resection and RPA abundance in FANCD2-depleted cells, but no decrease in the abundance of the RAD51-loading factor, BRCA2. Using QIBC, we found that FANCD2 and RAD51 colocalize at DSBs in G2 (**Figure S4G** – representative immunofluorescence images). Pairwise correlations between RAD51 and FANCD2 indicate that RAD51 and FANCD2 moderately colocalize in G2 (**Figure S4G** – plot). We therefore conclude that a subset of FANCD2 not only acts during DNA resection to alter DSB maturation, but also remains bound throughout RAD51-mediated recombination.

Finally, depletion of FANCD2 also alters repair protein recruitment to open chromatin sites in the genome. FANCD2 depletion causes an increase in MRE11, BLM, BRCA1, and BRCA2 at sites of open chromatin; but not RPA and RAD51 recruitment to these sites (**Figure S5A**). This observation suggests a broader role for FANCD2-FANCI in maintaining overall genome fidelity. The increase in the binding of other repair factors in the absence of FANCD2 could indicate that these sites experience increased DNA damage and mature into DSBs. As loading of FANCD2-FANCI to open chromatin is not ATM-dependent (**Figure 4D**), open chromatin sites may be under separate regulation, which would explain why these sites experience differential recruitment in the absence of FANCD2 than AsiSI DSBs. Overall, these results support a model in which FANCD2-FANCI are an integral part of the repair machinery that assembles onto open chromatin, stabilizing the structure and facilitating enzymatic activities of other DNA repair proteins.

### Chromatin-loaded FANCD2 promotes an ATR-dependent G2 cell cycle arrest

We showed that FANCD2-FANCI is bound to chromatin during the S and G2 cell cycle phases, removed from chromatin in mitosis, and excluded from chromatin in G1 (**Figure 3E** – plus extraction). Additionally, QIBC experiments revealed that DSB-dependent colocalization of FANCD2 with the HR repair factors BRCA1, RPA, and RAD51 occurred in G2 (**Figure 5E, S4G**). Therefore, we sought to understand whether chromatin-loaded FANCD2 itself could influence cell cycle progression.

Previous research has shown that >20 DSBs causes a G2 arrest, through activation of the G2/M DNA damage checkpoint^40^. Similarly, we found that induction of >100 DSBs using the AsiSI endonuclease causes asynchronously dividing cells to accumulate in G2 starting 4 hours after DSB induction and peaking around 20 hours after DSB induction (**Figure 6A** – siNTC). Depletion of FANCD2 using siRNA reduces the frequency of G2 cells (**Figure 6A** – siFANCD2). We confirmed that FANCD2-depleted cells have a reduced G2 accumulation 24 hours after DSB induction (**Figure 6B** – plus 4OHT / 24h). This reduction requires the FA core complex, as FANCA depletion by siRNA also reduces the G2 arrest 24 hours after DSB induction (**Figure S6A**). This observation of core dependency leads us to conclude that ubiquitinated FANCD2 clamped onto chromatin promotes an elongated G2 cell cycle arrest following DSB induction.

**Figure 6:**
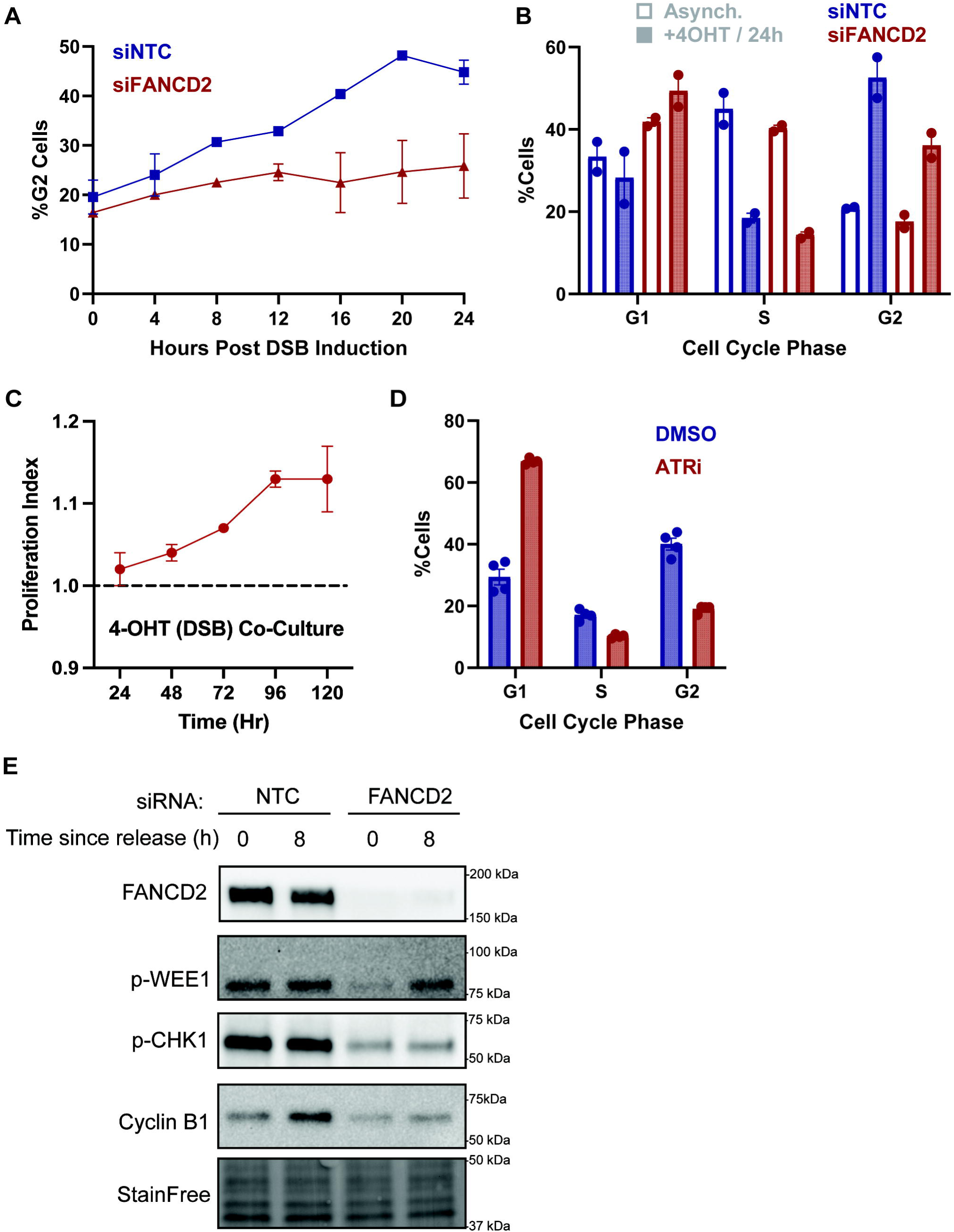
Chromatin-loaded FANCD2 promotes an ATR-dependent G2 arrest. A. Flow cytometry measurement of G2 abundance in mock (siNTC) or FANCD2-depleted (siFANCD2) cells at the indicated times after DSB induction. Data shown were generated from n=2 biological replicates. B. Flow cytometry cell cycle distribution of asynchronous (hollow plots) or 24-hour DSB-induced cultures (solid plots) in mock (siNTC) or FANCD2-depleted (siFANCD2) cells. Data shown were generated from n=2 biological replicates. C. Co-culture experiments comparing growth rates of CRISPRi non-targeting control (WT) and FANCD2 knockdown (fd2-) cells continuously grown in 4-OHT (which induces DSBs). Data shown are generated from n=2 biological replicates. D. Cell cycle distribution measured by QIBC of untreated or ATRi-treated cells 24 hours after DSB induction. Data shown are generated from at least m=12 images and n=2000 cells. E. Western blots showing phospho-CHK1 and phospho-WEE1 signal at indicated times after release from a double thymidine block into media containing 4-OHT in mock (siNTC) or FANCD2-depleted (siFANCD2) cells. Data shown are representative of n=3 blots.

The lack of a G2 accumulation in FANCD2-depleted cells and the corresponding increase in G1 cells may indicate that FANCD2-depleted cells undergo mitosis despite the presence of unrepaired DNA damage. We therefore hypothesized that FANCD2-depleted cells would divide more quickly in the presence of DNA damage than control cells. To test this hypothesis, we performed a co-culture assay using mock-treated and FANCD2-depleted cells. We found that FANCD2-depleted cells divided faster than the non-targeting controls when cultured in DSB-inducing conditions (**Figure 6C**). This data indicates that FANCD2-depleted cells not only continue to proliferate in the presence of DNA damage, but in fact proliferate more than control cells in the presence of DNA damaging agents. To validate this observation, we performed dye dilution flow cytometry experiments, in which patient-derived FA fibroblasts were pulse-labeled with a stable fluorescent dye. Dilution of this dye by proliferation allowed for cytometry-based tracking of cell proliferation. We observed that the FANCD2 null fibroblasts lost fluorescence more quickly than FANCD2-complemented cells in the presence of AluGG DSBs (**Figure S6B**). Based on the results of both the co-culture and dye dilution experiments, we conclude that FANCD2-deficient cells have less of a cell cycle delay under DSB-inducing conditions than wildtype cells. Accelerated cell division in the presence of DNA damaging agents appears to be a general feature of FA mutant cells, as shown by FANCD2-deficient cells proliferating faster than control cells in the presence of the crosslinking agent, MMC (**Figure S6C**).

DSB-induced G2 cell cycle arrest is known to be driven by ATR kinase^41^. Accordingly, inhibition of ATR using AZ20^42^ abrogates the G2 arrest in our cell lines (**Figure 6D**). Therefore, we explored links between FANCD2 and ATR signaling. RPA-coated single-stranded DNA is the main trigger for the recruitment and activation of ATR, with the help of 3 key ATR activator proteins (ATRIP, ETAA1, and TopBP1)^43^. Since ATR inhibition does not alter FANCD2 recruitment to DSBs 4 hours after DSB induction (**Figure S3B**), we speculate that the reduction in RPA loading at DSBs in FANCD2-depleted cells limits the recruitment and activation of ATR. Consistent with a model in which FANCD2 acts upstream of ATR activation, we observed that depletion of ATR activators reduced recombination in the DR-GFP system at an equivalent magnitude to FANCD2 depletion (**Figure S6D**).

ATR is known to drive the G2 cell cycle arrest through the ATR-CHK1-WEE1 axis^44^. Thus, one readout for ATR kinase activity is the accumulation of phosphorylated CHK1 and WEE1. We found that FANCD2-depleted cells accumulate less pCHK1 and pWEE1 when released from G1 synchronization in the presence of DSBs (**Figure 6E, S6E**). Phosphorylated WEE1 kinase inactivates CDK1 by phosphorylation, keeping the CDK-Cyclin B1 complex inactive and therefore maintaining a G2 arrest by preventing cells from entering mitosis^45^. Thus, less pWEE1 indicates that FANCD2-depleted cells do not arrest as strongly in G2 as control cells. Consistent with this result, immediately after release from the double thymidine block (0H), both control cells and FANCD2-depleted cells have similar Cyclin B1 levels. After 8H of DSB induction post release from the double thymidine block, control cells have accumulated high levels of Cyclin B1 as well as pWEE1, maintaining the cells in G2 due to high pWEE1 levels keeping CDK-Cyclin B1 inactive. In contrast, FANCD2-depleted cells have relatively low levels of Cyclin B1, which is consistent with the cells dividing, as Cyclin B1 is degraded during mitosis. We speculate that the reduction in RPA loading surrounding DSBs in FANCD2-deficient cells limits ATR recruitment, activation, and enforcement of a G2 arrest, which can be observed by the reduction in levels of pCHK1, a key substrate of ATR kinase. Altogether, these results indicate that FANCD2-FANCI both promotes DSB repair activities and contributes to cell cycle regulation to ensure that repair is completed before mitosis.

## Discussion

Here we report that FANCD2-FANCI functions as a chromatin surveillance complex that samples open chromatin regions and is subsequently stabilized at sites of DNA damage. FANCD2-FANCI efficiently localizes to transient open chromatin formed at DSBs in a manner that is dependent on the FA core complex and ATM kinase activity. Chromatin-loaded FANCD2-FANCI maintains chromatin accessibility and stabilizes BRCA1 and BLM at DSBs, thereby increasing DSB resection and RPA loading. FANCD2-FANCI also contributes to a G2 arrest in response to DNA damage. Together, these FANCD2-FANCI activities promote resection-based DSB repair outcomes.

Reframing the FANCD2-FANCI heterodimer as a chromatin surveillance factor presents a unifying explanation for its diverse substrates and activities. There are many open chromatin sites within cells that have been reported to interact with FANCD2-FANCI, including mitochondrial DNA^27^, extrachromosomal DNA^28^, viral DNA^46^, transcriptional start sites^47^, and genomic DNA regulatory elements^48^. In addition, FANCD2-FANCI has been reported to coordinate repair events at DNA lesions, which form transient open chromatin regions. We therefore propose that the primary role of FANCD2-FANCI is to associate with open chromatin and coordinate DNA repair activities in response to local regulation. These regulatory signals include ATM at DSBs and ATR at stalled replication forks. Our results establish specific roles for FANCD2-FANCI in promoting resection and chromatin openness at DSBs. However, there are likely to be additional regulatory signals and scaffolding roles that remain to be characterized at other sites of FANCD2-FANCI loading.

We and others have previously demonstrated that FANCD2-FANCI promotes DSB repair by HR^12,13,49^. Here we use inducible targeted nucleases to investigate repair events at approximately 200 distinct DSB sites in the genome. Our results show that FANCD2-FANCI promotes resection and RPA loading at DSBs. When compiling our stranded ChIP-seq data, we were surprised by the number of DSB sites that loaded RPA and RAD51. This observation overturns assumptions about the limiting step in HR. The dominant model is that DSB resection occurs rarely and as a precursor to recombination. Our results support a model in which resection, and even loading of RAD51, is common at DSBs in proliferating cells. This finding suggests that fill-in synthesis or recombination using a sister chromatid or homolog as a template is far more common than hypothesized. To this latter point, we note that there is significant RPA loading on the strand containing the 5’ phosphate on either side of the DSB (**Figure 5D**). This signal has previously been shown to arise from rebinding of RPA to the displaced strand during strand exchange in meiotic recombination^50^. We therefore speculate that the rate-limiting step in HR repair of DSBs in proliferating cells occurs after strand exchange between the DSB and the template.

Our ChIP-seq and DSB repair results suggest that FANCD2-FANCI cooperates with BRCA1 but not BRCA2 during DNA repair. The ChIP-seq distribution of FANCD2-FANCI on chromatin matched the distribution of BRCA1/RPA/BLM but differed from the distribution of MRE11/BRCA2. This result is consistent with a model in which the FANCD2-FANCI heterodimer cooperates with BRCA1 and BLM as part of a larger protein assembly to promote DSB resection, and is distinct from the roles of MRE11 and BRCA2 in this process. Our knockdown experiments further support a model in which FANCD2-FANCI and BRCA1 have a shared function in promoting resection that is distinct from the role of BRCA2. Individual knockdown of FANCD2 and BRCA1 decreases DR-GFP recombination (**Figure S4B**), while dual knockdown of both FANCD2 and BRCA1 reduces recombination still further. On the other hand, individual knockdown of BRCA2 decreases recombination, but there is no additive effect during dual knockdown of FANCD2 and BRCA2 (**Figure S4B**). One possibility to explain these observations is that loaded FANCD2-FANCI acts as an anchor to stabilize other DNA repair factors during DSB resection.

The wide range of DNA substrates bound to FANCD2-FANCI supports persistent association of the complex with chromatin during DSB repair. We and others have shown that FANCD2 binds single-stranded and double-stranded DNA spanning the cut site (**Figure 1G**). We have observed DNA spanning the cut site with deletions characteristic of EJ outcomes (**Figure S1A**)^51^, as well as single-stranded DNA ending at the cut site (**Figure 1F**). These outcomes are consistent with different roles for FANCD2-FANCI: chromatin maintenance, EJ DSB repair, and HR DSB repair, respectively. To reconcile these observations, we propose that FANCD2-FANCI interacts wherever naked DNA is exposed between DNA-bound protein complexes. Signaling from a second complex – ATM in proximity to DSBs – triggers ubiquitination and clamping of FANCD2-FANCI onto chromatin. This loaded form of FANCD2-FANCI acts as a scaffold for other repair factors and remains loaded throughout DNA repair processes.

This model has two unique features. First, as FANCD2-FANCI binds to open chromatin and not the lesion itself, resolution of the lesion does not require removal of FANCD2-FANCI. Second, encirclement of DNA by ubiquitinated FANCD2-FANCI permits association of the heterodimer throughout remodeling events at the DSB. We speculate that this binding model allows persistence of FANCD2-FANCI on chromatin and explains why we see multiple DNA substrates bound to FANCD2-FANCI using bulk ChIP-based approaches. We note that FANCD2 loads to DSBs after MRE11 (**Figure 4A**), but DSBs could be resolved by EJ prior to long-range resection. If EJ occurs, it will produce FANCD2-FANCI bound to intact or deletion-modified double-stranded DNA. In other cases, FANCD2-FANCI can facilitate DSB resection, which produces FANCD2-FANCI bound to single-stranded DNA. Future studies will be needed to define the precise conditions that allow the unloading of FANCD2-FANCI from chromatin; however, our results indicate that bulk unloading of FANCD2-FANCI occurs before the completion of mitosis, and thus may be a coordinated and active process.

FANCD2-FANCI loading onto chromatin further regulates cell cycle progression after DSB induction. We observed that FANCD2-depleted cells have a reduced G2 accumulation compared to control cells (**Figure 6A, 6B**). The lack of a G2 accumulation in FANCD2-depleted cells, and the corresponding increase in G1 cells, could indicate that FANCD2-depleted cells either divide despite the presence of DNA damage or repair DSBs faster using ligation-based EJ repair. Our results support a model in which the absence of chromatin-loaded FANCD2-FANCI reduces DSB processing (by resection), which in turn reduces DSB repair time. This shorter repair duration requires a shorter G2 arrest despite the presence of many DSBs. We note that DSB-damaged control cells accumulated in G2 until 20 hours post induction, while DSB-damaged FANCD2-depleted cells reached peak G2 accumulation approximately 12 hours after DSB induction (**Figure 6A**). Since FANCD2 promotes DNA resection and RPA loading onto single-stranded DNA, FANCD2 is upstream of the ATR-CHK1-WEE1 signaling cascade, which prevents mitotic entry in the presence of DNA damage.

Our data further demonstrate that FANCD2-FANCI chromatin association is cell cycle regulated, occurring primarily during S and G2 phases. The cell cycle dependent activity of FANCD2-FANCI allows the heterodimer to coordinate various DNA repair activities in response to unique regulatory signals. During S phase, ATR activates the heterodimer to act at stalled replication forks as part of replication-associated repair^38^. During G2 phase, ATM activates the heterodimer to act at DSBs as part of template-dependent HR mechanisms. Its exclusion from chromatin during G1 reinforces its specialized function in facilitating template-dependent HR mechanisms at DSBs, as the sister chromatid is available to serve as the template molecule specifically during G2. Interestingly, FANCD2-FANCI was also found on mitochondrial DNA and extrachromosomal DNA (**Figure S2A**), suggesting that its chromatin surveillance functions are not limited to the nuclear genome.

Finally, FANCD2-FANCI’s binding to open chromatin suggests it also regulates undamaged genomic regions. A major prediction of this surveillance model is that large amounts of DNA damage could sequester FANCD2-FANCI away from normally open sites, thereby preventing its usual function in stabilizing these regions. Our data show that FANCD2-FANCI stabilizes open chromatin sites and prevents their progression into MRE11-bound DSBs (**Figure S5A**). Accordingly, any reduction – direct or indirect – in FA protein levels may elevate spontaneous lesions at open chromatin sites, increasing genomic stress. In this sense, FA proteins serve as both proactive sensors, preserving genome fidelity by preventing endogenous damage, and reactive effectors, repairing exogenous DNA insults. Taken together, these findings support a unifying model in which the FA-BRCA pathway acts as a sentinel to maintain genomic integrity.

## Acknowledgements

We thank Gaëlle Legube (Center for Integrative Biology, Toulouse, France) for DIvA AsiSI-ER-U2OS cells and DIvA-AID AsiSI-ER-U2OS cells. We thank Jeremy Stark (City of Hope, Duarte, California, USA) for U2OS DR-GFP cells. We thank Leslie Wakefield and the Fanconi Anemia Research Materials (FARM) repository (located at Oregon Health & Science University and funded by the Fanconi Cancer Foundation) for providing Fanconi anemia patient-derived fibroblast cell lines. We thank the members of the Gardner and Richardson labs for their feedback on the manuscript. We acknowledge the assistance of Dr. Jennifer Smith, manager of the Biological Nanostructures Laboratory within the California NanoSystems Institute, supported by the University of California, Santa Barbara and the University of California, Office of the President. C.D. Richardson acknowledges support from National Institutes of Health R35GM142975. B.M. Gardner acknowledges support from National Institutes of Health K99/R00GM121880, R35GM146784, and the Searle Scholars Program. C.M. Joyce acknowledges support from the Jane Altman Fellowship. S.P. Chowdhury acknowledges support from the Storke Family Fellowship. A.K. Wang and Z.N. Rodriguez acknowledge support from the MARC Scholars Program. C. Cruz acknowledges support from an administrative supplement to R35GM142975. Y. Tzadikario acknowledges support from the CIRM COMPASS Program. The content is solely the responsibility of the authors and does not necessarily represent the official views of the National Institutes of Health.

## Author contributions

C.M. Joyce: Conceptualization, Formal analysis, Investigation, Methodology, Validation, Visualization, Writing – original draft, Writing – review and editing.

J. Bacal: Investigation, Methodology, Validation, Visualization.

S.P. Chowdhury: Formal analysis, Investigation, Methodology, Validation, Visualization, Writing – Review and editing.

A.N. Brown: Investigation, Validation.

A.K. Wang: Investigation, Validation.

C. Cruz: Investigation, Validation.

K. Bains: Investigation.

Z.N. Rodriguez: Investigation.

N.J. McCormick: Investigation.

Y. Tzadikario: Investigation.

K.U. Tavasoli: Investigation.

B.M. Gardner: Conceptualization, Methodology, Investigation, Supervision, Writing – review & editing.

C.D. Richardson: Conceptualization, Data curation, Formal analysis, Funding acquisition, Investigation, Methodology, Project administration, Software, Resources, Supervision, Validation, Visualization, Writing – original draft, Writing – review & editing.

## Declaration of interests

None

## Data Accessibility

Raw data for ChIP and ATACseq have been uploaded to SRA under PRJNA1249941. Kas-ATAC seq data from GM16828 cells was reprocessed from PRJNA1103095. Induced and uninduced γ-H2AX ChIP-seq reads in DIvA cells were reproduced from PRJEB21297. DNase-seq in MG63 cells was reprocessed from ENCODE ENCFF790EMB. Repli-seq data in U2OS cells was reprocessed from PRJNA397123.

---

**Figure S1:**
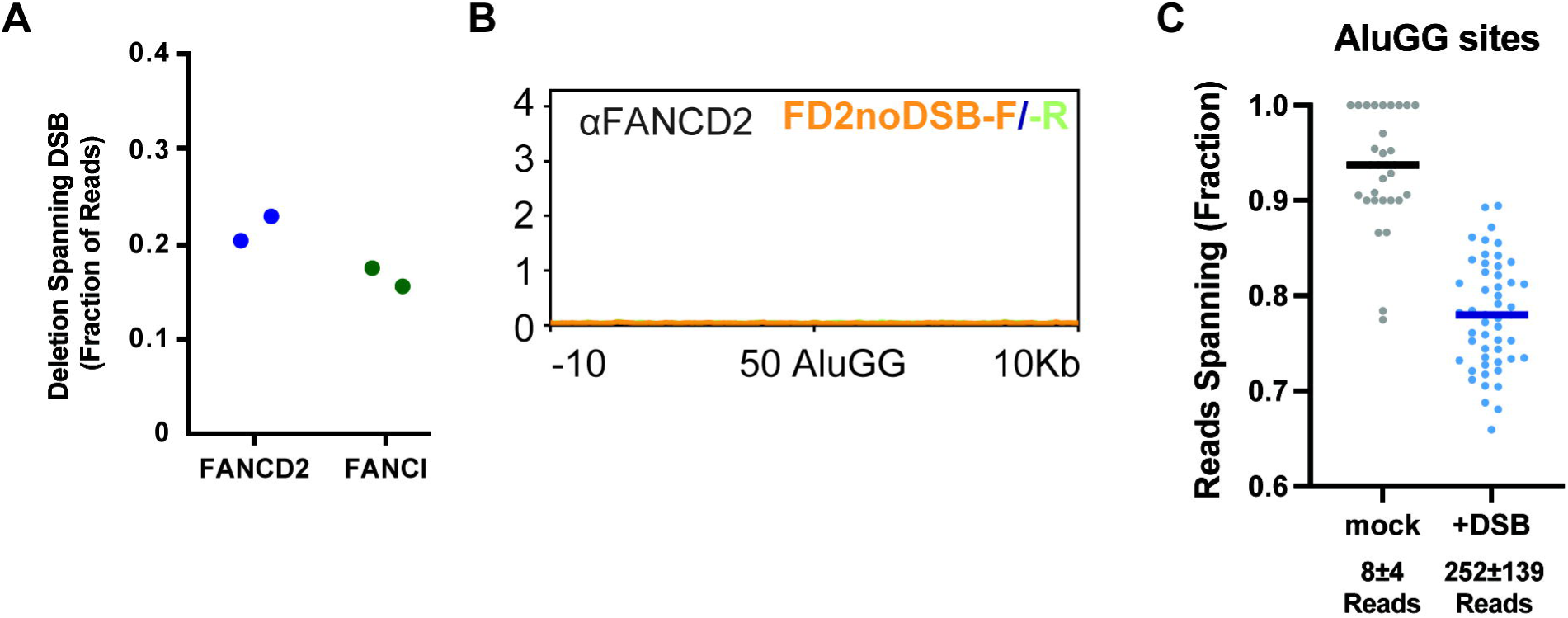
A. Frequency of ChIP reads within a 200bp window surrounding the HBB DSB that encode a 9bp deletion that spans the site of the Cas9 DSB in FANCD2 (blue) or FANCI (green) ChIP-seq experiments. Data from n=2 biological replicates is plotted. B. Stranded ChIP-seq data presented as reads mapping to distinct strands at 50 AluGG sites in their natural context (no DSB). Immunoprecipitations were performed using a FANCD2 antibody. All plots are representative of n=2 biological replicates. C. Frequency of individual reads in the FANCD2 ChIP dataset from Figure 1D spanning the AluGG DSB sites. All data from n=2 biological replicates is plotted.

**Figure S2:**
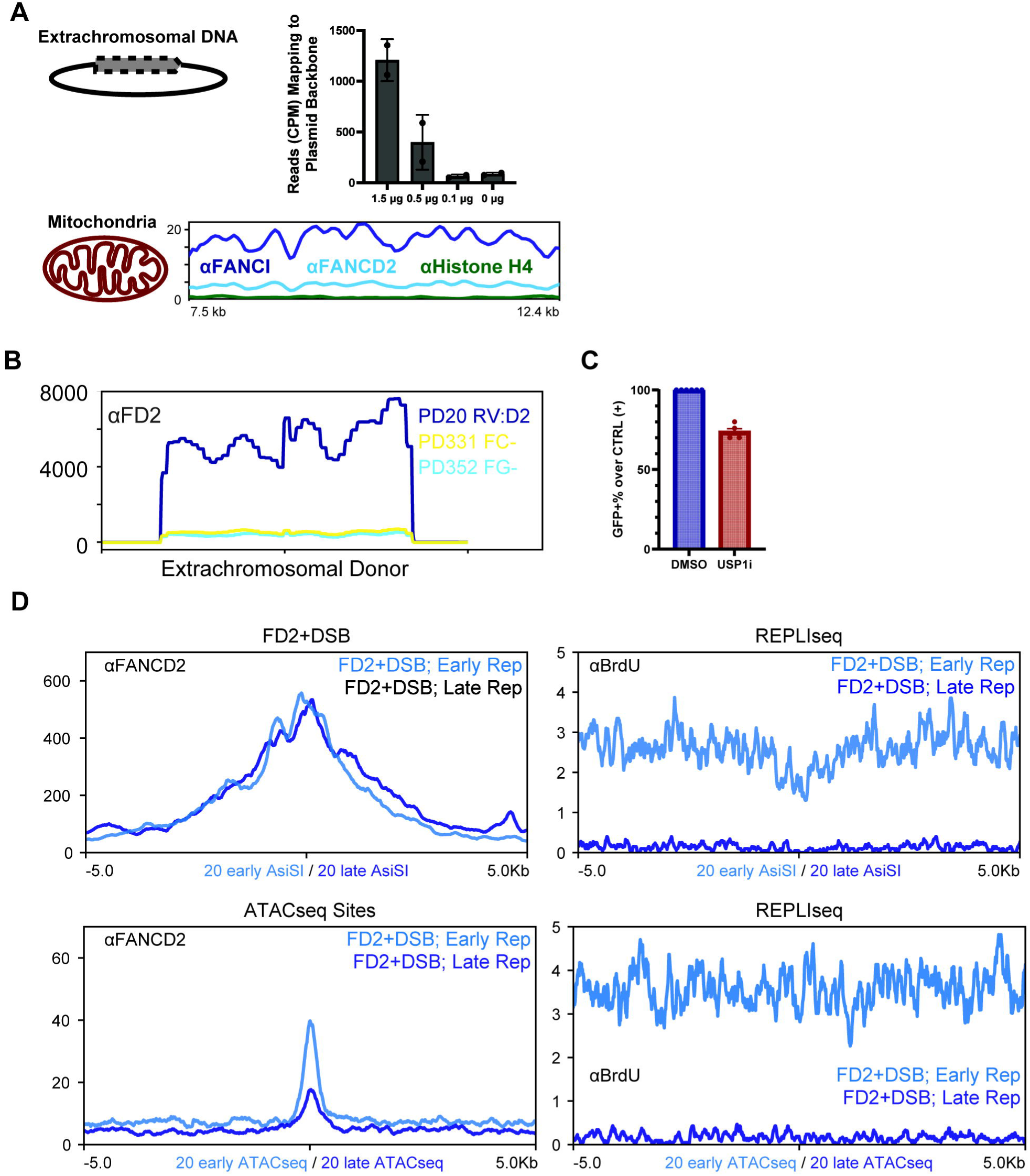
A. (Top) Total FANCD2 ChIP-seq reads (in counts per million) aligning to the pMB1-ampR plasmid backbone region of extrachromosomal DNA homologous to the HIST1H2BJ or LMNB1 locus in K562 cells electroporated with Cas9 RNP targeting the HIST1H2BJ locus. (Bottom) ChIP-seq profiles for FANCD2, FANCI, and Histone H4 binding to mitochondrial DNA. Data shown are generated from n=2 biological replicates. B. ChIP-seq data showing FANCD2 binding to extrachromosomal template DNA in cell lines expressing FANCD2 (PD20 RV:D2), or lacking the FA core components FANCC (PD331) or FANCG (PD352). Data shown are representative of n=2 biological replicates. C. Normalized recombination (GFP+) percentage at the DR-GFP locus in cells either mock-treated (DMSO) or USP1 inhibitor-treated (USP1i) cells. Data generated from at least n=3 biological replicates. D. ChIP-seq (left) and REPLIseq (right) data at AsiSI DSB and ATAC-seq open chromatin sites. REPLIseq data is reproduced from PRJNA397123. Data shown are representative of n=2 biological replicates.

**Figure S3:**
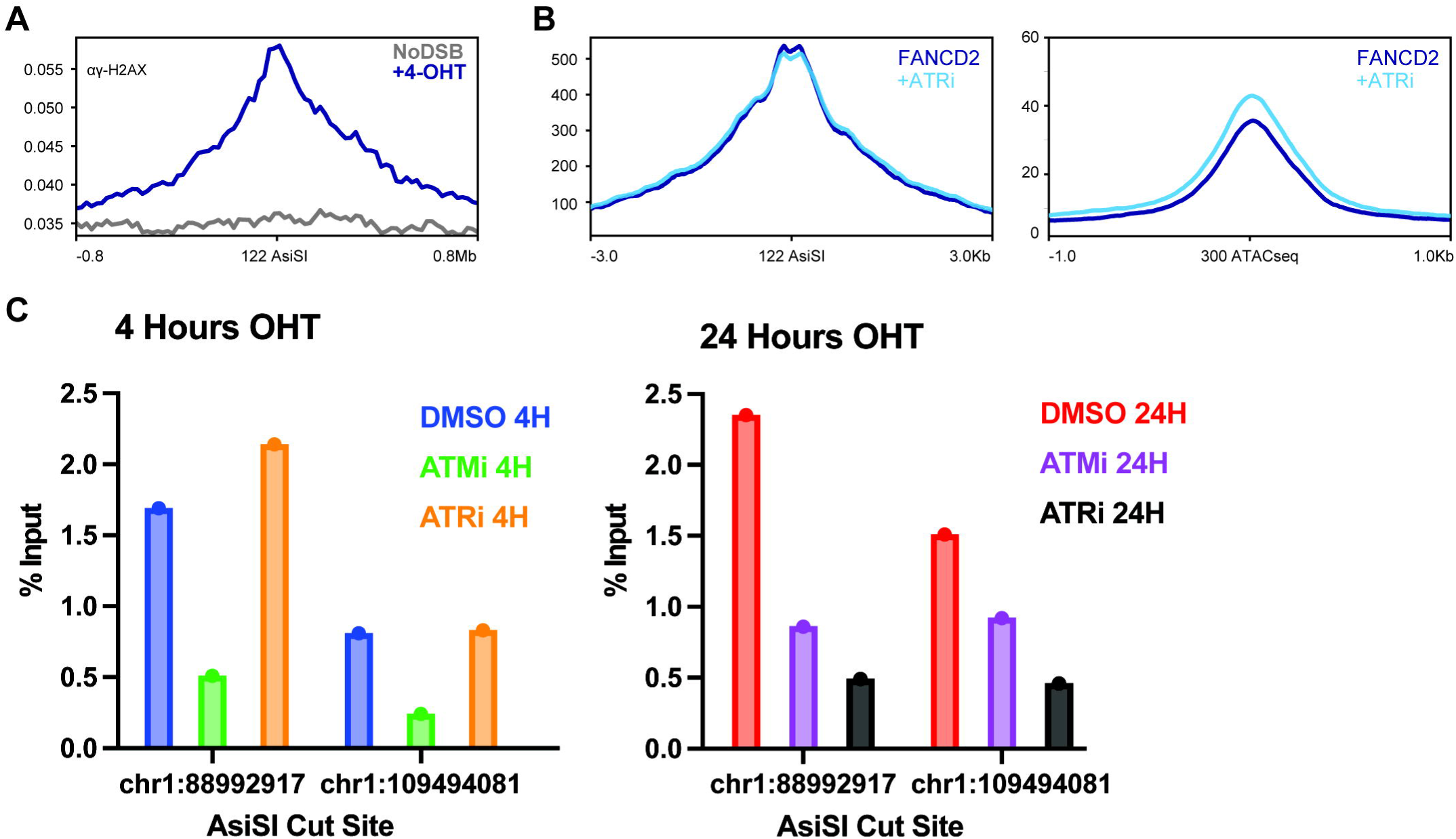
A. ChIP-seq data showing γ-H2AX binding to AsiSI sites in U2OS cells after DSB induction. Data shown is reproduced from PRJEB21297 and representative of n=2 biological replicates. B. ChIP-seq summary showing FANCD2 binding to AsiSI DSBs (left) or open chromatin sites (right) in untreated, or AZ20-treated conditions. Data shown are representative of n=2 biological replicates. C. ChIP-qPCR data showing FANCD2 enrichment at two AsiSI DSB sites in the presence of ATM (KU-55933) or ATR (AZ20) inhibitors 4 (left) or 24 (right) hours after DSB induction.

**Figure S4:**
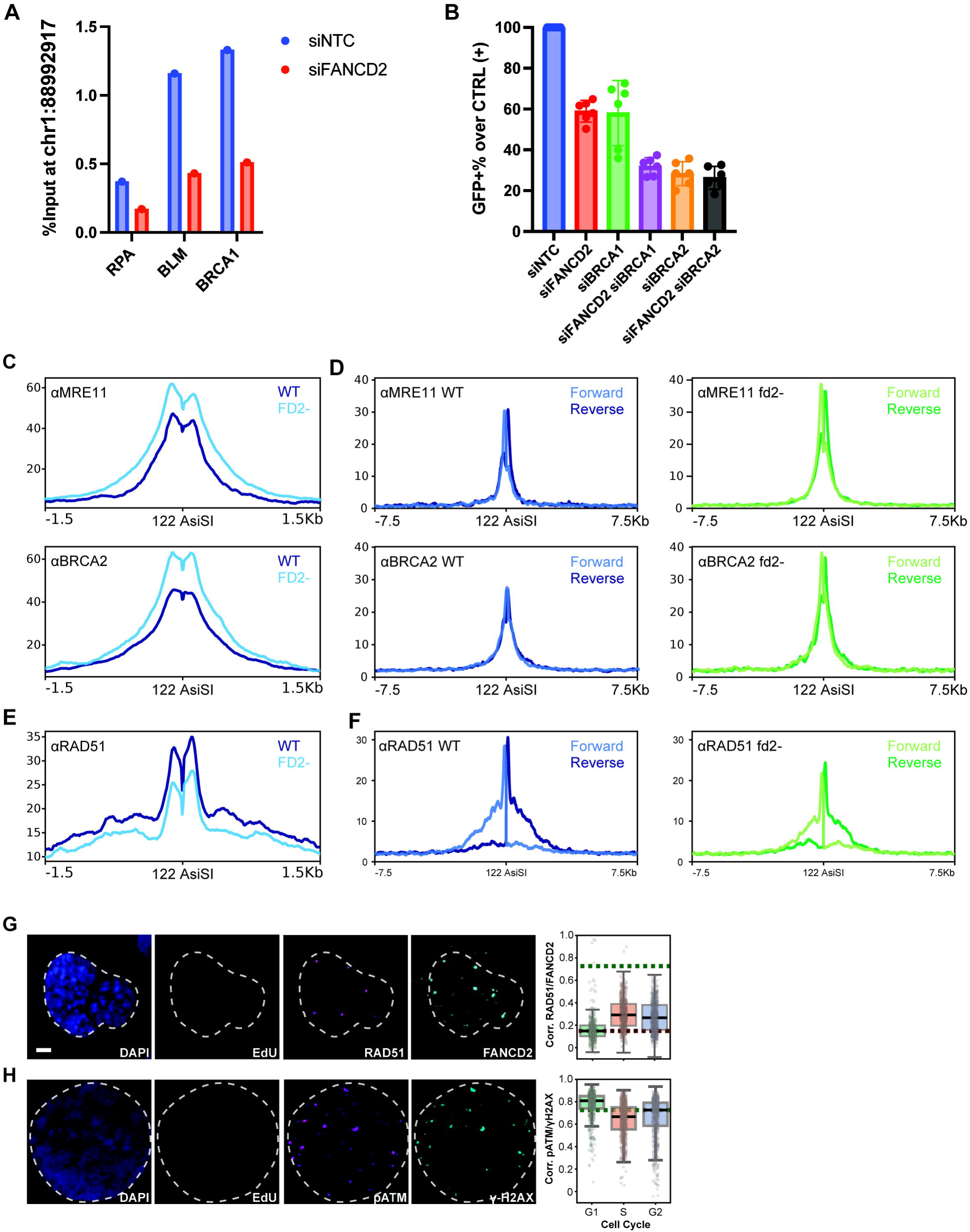
A. ChIP-qPCR data showing recruitment of RPA, BLM, and BRCA1 to a single AsiSI cut site 4 hours after DSB induction in WT (siNTC) or FANCD2-depleted (siFANCD2) cells. Data shown were generated from n=3 technical replicates. B. Normalized recombination (GFP+) percentage at the DR-GFP locus in cells with the indicated siRNA knockdowns. Data generated from at least n=3 biological replicates. C. ChIP-seq data showing recruitment of MRE11 or BRCA2 to AsiSI DSBs in CRISPRi non-targeting control (WT) and FANCD2 knockdown (fd2-) cells 4 hours after DSB induction. Data shown are representative of n=2 biological replicates. D. Stranded ChIP-seq data showing recruitment of MRE11 or BRCA2 to AsiSI DSBs in CRISPRi non-targeting control (WT) and FANCD2 knockdown (fd2-) cells 4 hours after DSB induction. Data shown are representative of n=2 biological replicates. E. ChIP-seq data showing recruitment of RAD51 to AsiSI DSBs in CRISPRi non-targeting control (WT) and FANCD2 knockdown (fd2-) cells 4 hours after DSB induction. Data shown are representative of n=2 biological replicates. F. Stranded ChIP-seq data showing recruitment of RAD51 to AsiSI DSBs in CRISPRi non-targeting control (WT) and FANCD2 knockdown (fd2-) cells 4 hours after DSB induction. Data shown are representative of n=2 biological replicates. G. QIBC data showing colocalization of DAPI, EdU, and RAD51 with FANCD2. Example images are presented at left and correlation is plotted at right. Data shown are generated from at least m=12 images and n=2000 cells. Scale bar = 2μm. H. QIBC data showing colocalization of DAPI, EdU, and phospho-ATM with γ-H2AX. Example images are presented at left and correlation is plotted at right. Data shown are generated from at least m=12 images and n=2000 cells.

**Figure S5:**
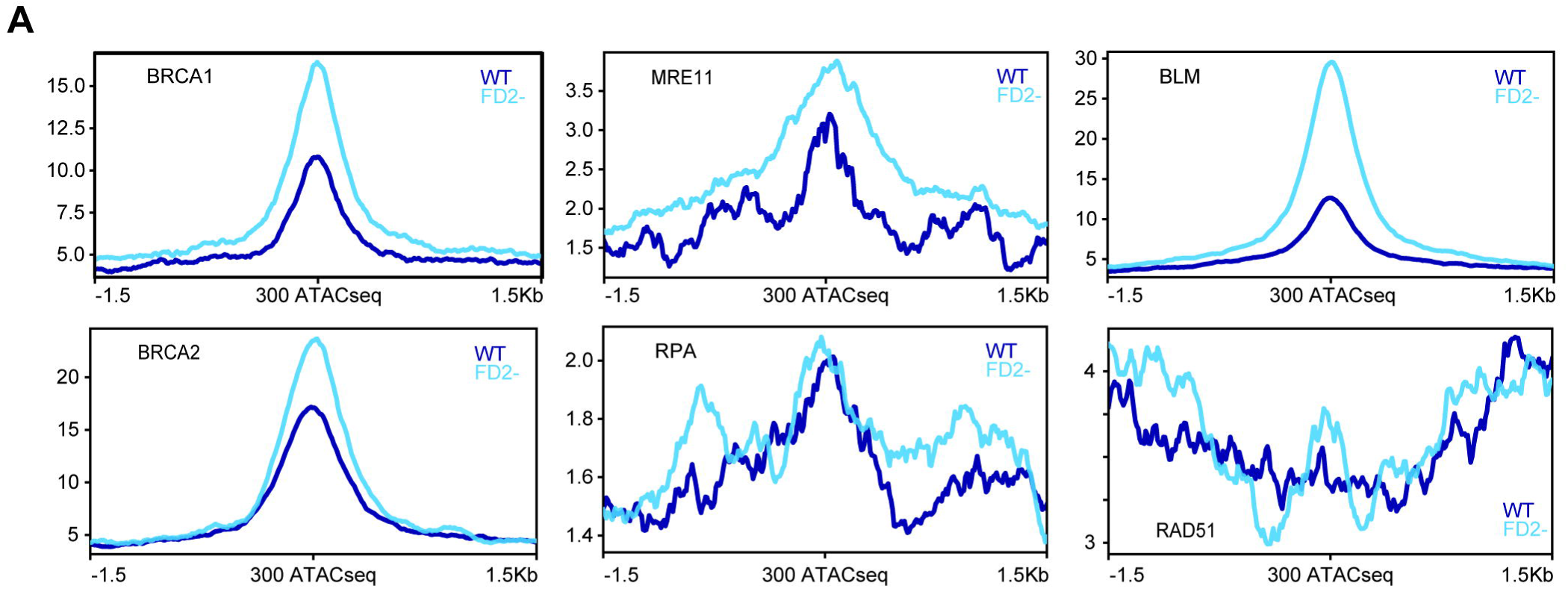
A. ChIP-seq data showing recruitment of BRCA1, MRE11, BLM, BRCA2, RPA, or RAD51 to open chromatin sites in CRISPRi non-targeting control (WT) and FANCD2 knockdown (fd2-) cells. Data shown are representative of n=2 biological replicates.

**Figure S6:**
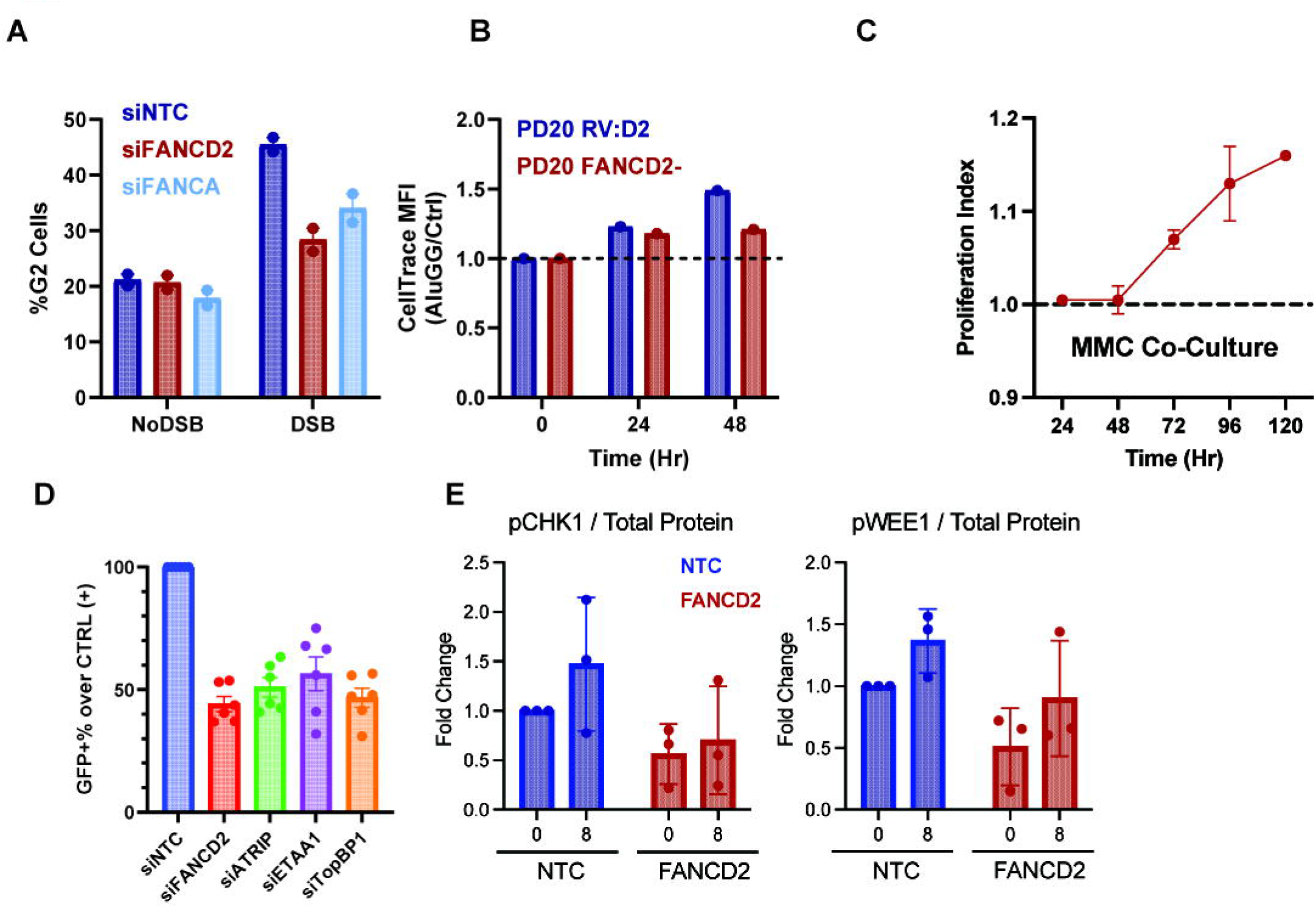
A. QIBC measurement of G2 abundance in mock (siNTC), FANCD2-depleted (siFANCD2), or FANCA-depleted (siFANCA) cells after 24 hours of mock (noDSB) or 4-OHT (DSB) treatment. Data shown are generated from at least m=12 images and n=2000 cells. B. CellTrace signal normalized to control (no DSB) condition at indicated timepoints after DSB induction in WT (PD20 RV:D2) or FANCD2^-/-^ (PD20) cells. Data shown are generated from n=1 biological replicates. C. Co-culture experiments comparing growth rates of CRISPRi non-targeting control (WT) and FANCD2 knockdown (fd2-) cells continuously grown in MMC (which induces ICLs). Data shown are generated from n=2 biological replicates. D. Normalized recombination (GFP+) percentage at the DR-GFP locus in cells with the indicated siRNA knockdowns. Data generated from at least n=3 biological replicates. E. Quantification of Western blots shown in Figure 6E.

## Notes

### Competing Interest Statement

The authors have declared no competing interest.

